# Lagrangian Formalism in Biology: II. Non-Standard and Null Lagrangians for Nonlinear Dynamical Systems and their Role in Population Dynamics

**DOI:** 10.1101/2023.01.18.524517

**Authors:** D. T. Pham, Z. E. Musielak

## Abstract

Non-standard Lagrangians do not display any discernible energy-like terms, yet they give the same equations of motion as standard Lagrangians, which have easily identifiable energy-like terms. A new method to derive non-standard Lagrangians for second-order nonlinear differential equations with damping is developed and the limitations of this method are explored. It is shown that the limitations do not exist only for those nonlinear dynamical systems that can be converted into linear ones. The obtained results are applied to selected population dynamics models for which non-standard Lagrangians and their corresponding null Lagrangians and gauge functions are derived, and their roles in the population dynamics are discussed.

## 1 Introduction

In physics equations of motion are derived by using the Lagrangian formalism, which requires knowledge of a function called Lagrangian. By specifying a Lagrangian for a given system, its equation of motion is derived by substituting this Lagrangian into the Euler-Lagrange (E–L) equation. There are three different families of Lagrangians, namely, standard, non-standard, and null.

Standard Lagrangians represent the difference between the kinetic and potential energy [1], and they play a central role in Classical Mechanics [2–4] as well as other areas of physics [5]. Non-standard Lagrangians (NSLs) are different in form from standard Lagrangians but also give the same equation of motion as the NSLs. The main difference is that NSLs lack terms that clearly discern energy-like forms [6]. There are also null Lagrangians and they identically satisfy the E-L equation and can be expressed as the total derivative of any scalar function [7]. Since null Lagrangians have no effect on equations of motion, they are not discussed any further in this paper.

Different methods to derive standard [8–15] and nonstandard [12–26] Lagrangians were developed. The methods were used to obtain both types of Lagrangians for many different physical systems. In the first attempts to obtain Lagrangians for selected biological systems, the Lagrangians were found by guessing their forms [27–29]. Then, they were derived using the Method of Jacobi Last Multiplier [30,31]. More recently, the first standard Lagrangians were derived for five population dynamics models and their implications for the models were discussed [32].

This paper aims to extend this work by deriving NSLs for selected population dynamics models considered in [32], which are: the Lotka-Volterra [33,34], Verhulst [35], Gompertz [36], Host-Parasite [37], and SIR [38] models. The models are described by second-order nonlinear and damped ODEs, for which a new method is required to find NSLs. In this paper, a novel method is developed and then used to derive NSLs for all considered population models. The derived NSLs are compared to those previously obtained [29,30,32] and used to gain new insights into the role they play in the models, and in the symmetries underlying them. For the derived NSLs, their corresponding null Lagrangians (NLs) and gauge functions are also calculated, and these NLs in the population dynamics is discussed.

The paper is organized as follows. Section 2 presents our method to construct non-standard Lagrangians and its limitations; in Section 3, the population dynamics models are described and our method to derive their non-standard Lagrangians for these models are presented; then, non-standard Lagrangians for the population dynamics models are given in Section 4; null Lagrangians and their gauge functions are derived and discussed in Section 5; the obtained results are compared to those previously obtained and discussed in Section 6; and our Conclusions are given in Section 7.

## 2 Lagrangian formalism

### 2.1 Action and non-standard Lagrangians

The functional 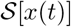 is called action and is defined by an integral over a scalar function *L* that depends on the differentiable function *x*(*t*) that describes the time evolution of any dynamical system, and on the time derivative 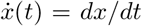. The function 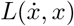 is called the Lagrangian function or simply *Lagrangian,* and it is a map from the tangent bundle 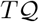 to the real line **R**, or 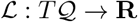, with 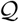 being a configuration manifold [3]. In general, the Lagrangian may also depend explicitly on *t*, which can be written as 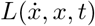 and requires 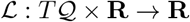.

According to the principle of least action, or Hamilton’s principle [2–6], the action 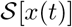 must obey the following requirement 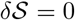, which guarantees that the action is stationary, or has either a minimum, maximum, or saddle point. The necessary condition that 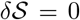, is known as the E-L equation, whose operator 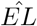 acts on the Lagrangian, 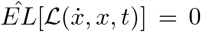 and gives a second-order ODE that becomes an equation of motion for a dynamical system with 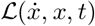. The process of deriving the equation of motion is called the Lagrangian formalism [2–4], and it has been extensively used in physics to derive its fundamental classical and quantum equations [5].

The Lagrangian formalism is valid for both standard and non-standard Lagrangians but only the latter are considered in this paper. There are different forms of NSLs [12–22] but only two of them are considered in this paper. The first considered form of NSL is

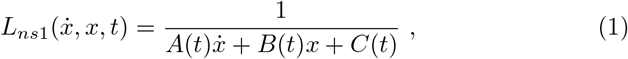

and it is applied to the equation of motion given by

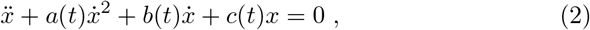

where the coefficients *a*(*t*), *b*(*t*) and *c*(*t*) are at least twice differentiable functions of the independent variable *t*, and the functions *A*(*t*), *B*(*t*) and *C*(*t*) are expressed in terms of these coefficients by substituting 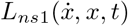 into the E–L equation.

The second form of the considered NSL is

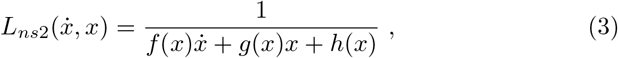

and it is used for the following equation of motion

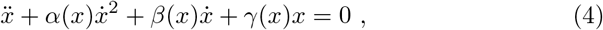

where the coefficients *α*(*x*), *β*(*x*) and *γ*(*x*) are at least twice differentiable functions of the dependent variable *x*(*t*), and the functions *f*(*x*), *g*(*x*) and *h*(*x*) are determined by substituting 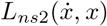 into the E–L equation, which allows expressing these functions in terms of the coefficients.

The NSL 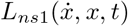 was extensively studied and applied to several dynamical systems [12,13,20]. However, studies of 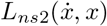 were limited to only some special cases [20]. Therefore, one of the objectives of this paper is to perform a detailed analysis of 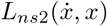 and its applicability to the population dynamics models; so far, only other forms of NSLs been applied to the models [27–31]. Moreover, in a recent work [32], the standard Lagrangians were constructed for the same sample of the population dynamics models.

### 2.2 Limits on construction of non-standard Lagrangians

To obtain a NSL means to determine the unknown functions in Eqs (1) and (3), which requires that an equation of motion is given. As shown in [32], the equations of motion for the population dynamics models can be written in the following form

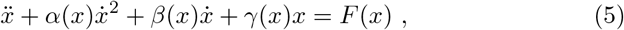

which generalizes Eq. (4) by taking into account a driving force *F*(*x*). For different models, the coefficients *α*(*x*), *β*(*x*) and *γ*(*x*) are given, and *F*(*x*) is also known. Since Eq. (5) does not depend explicitly on time, the Lagrangian that can be used for this equation is 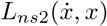 given by Eq. (3). From now on, we shall take 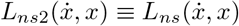 and use the latter throughout this paper.

It is seen that Eq. (5) has both linear and quadratic damping terms, that can become nonlinear depending on the form of *γ*(*x*)*x*, and that it is driven. In a previous study [32], the standard Lagrangian for Eq. (5) was found in case *F*(*x*) = const. To construct NSL for Eq. (5), we take 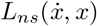 given by Eq. (3) and determine the functions *f*(*x*), *g*(*x*) and *h*(*x*) in terms of the known coefficients *α*(*x*), *β*(*x*) and *γ*(*x*). Our approach generalizes the previous work [18], and it allows us to investigate the applicability of 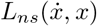 to Eq. (5) and its limitations.

#### Proposition 1

Let a general non-standard Lagrangian be given by

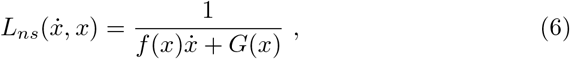

where *G*(*x*) = *g*(*x*)*x* + *h*(*x*). Then, 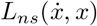 becomes the NSL for Eq. (5) if, and only if,

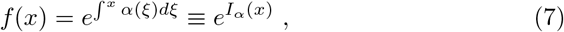

and

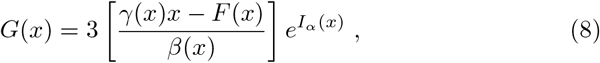

and either one of the following conditions

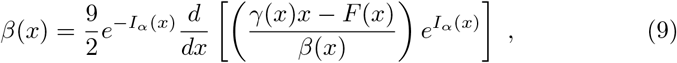

and

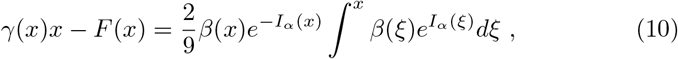

is satisfied.

#### Proof

Substituting Eq. (5) into the E–L equation, we obtain

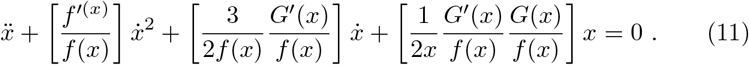

By comparing this equation to Eq. (5), we find the following relationships

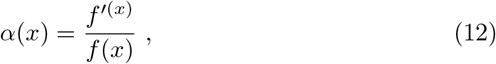

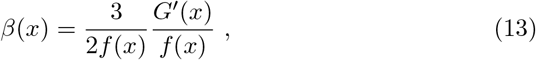

and

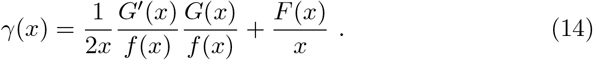

Using Eq. (12), we obtain *f*(*x*) given by Eq. (7). However, by combining Eqs (13) and (14), and using Eq. (12), we find *G*(*x*) given by Eq. (8). Then, the conditions expressed by Eqs (9) and (10) are easy to derive from Eqs (12), (13) and (14). Moreover, substituting Eq. (10) into Eq. (9) shows that both conditions are equivalent, thus, it is sufficient to use only one of them. This concludes the proof.

#### Corollary 1

The coefficients *β*(*x*) and *γ*(*x*) mutually depend on each other through the function *G*(*x*) and, in addition, they also depend on *α*(*x*) through the function *f*(*x*).

#### Corollary 2

The conditions given by Eqs (9) and (10) are equivalent, which means that if one of them is not satisfied then 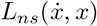, with *f*(*x*) and *G*(*x*) (see Eqs 7 and 8), cannot be considered to be the NSL for Eq. (5).

#### Corollary 3

Only *α*(*x*) is required to uniquely determine the function *f*(*x*).

### 2.3 From nonlinear to linear equations

The main result of Proposition 1 is that any NSL of the form of Eq. (6) can only be constructed if one of the conditions given by Eqs (9) and (10) is satisfied. We now explore other consequences of the condition given by Eq. (10) in the following proposition.

#### Proposition 2

Let the equation of motion be

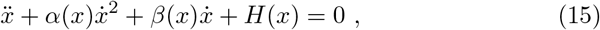

where *H*(*x*) = *γ*(*x*)*x* – *F*(*x*), and let

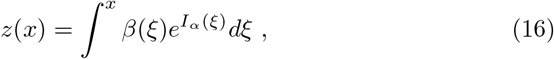

be a new dependent variable that can be expressed in terms of a new variable *η* that is related to the original variable *x* by *dη* = *β*(*x*)*dx*. Then, Eq. (15) becomes

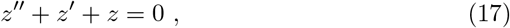

with *z*′ = *dz/dη* and *z*″ = *d*^2^*z/dη*^2^, if, and only if,

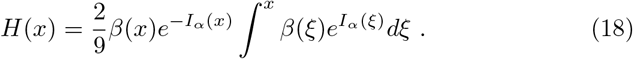

#### Proof

Using Eq. (16), we find

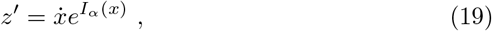

and

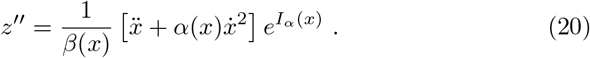

Substituting Eqs (19), (20) and (16) into Eq. (17), we obtain the original equation of motion (see Eq. 15) with the function *H*(*x*) given by Eq. (18). This concludes the proof.

#### Corollary 4

Equation (17) is the Sturm-Liouville equation [43] whose solutions are well-known and given as 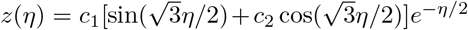.

#### Corollary 5

The non-standard Lagrangian given by Eq. (6) can be constructed without any limits for those nonlinear ODEs that can be converted into linear ODEs, which requires that the condition given by Eq. (10) is satisfied.

### 2.4 Non-standard Lagrangians without limits

The results of Proposition 2 and Corollaries 4 and 5 demonstrate that the existence of NSL of the form of Eq. (6) is determined by the conditions given by Eqs (9) and (10), which must be satisfied in order for the NSL to exist. According to Corollaries 1-3, the only two terms in the equation of motion given by Eq. (5) that can be uniquely derived without any limits by the NSL are the terms with 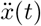 and 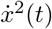. The coefficients in terms with 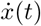 and *x*(*t*) are mutually related, thus, they are dependent on each other as shown by either Eq. (9) or Eq. (10).

Because of these limitations on the construction of NSLs for Eq. (5), let us now propose another method by extending the previous work [32]. The basic idea of this work is to write Eq. (5) in the following form

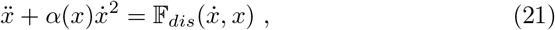

where

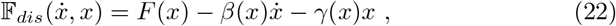

becomes a dissipative force because of its dependence on 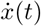. In this case, the E–L equation [2,3,32] can be written as

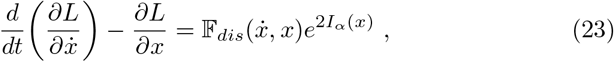

where the force on the RHS of this equation is known at the Rayleigh force [2,3]

Based on the results presented in Section 2.2, the NSL for Eq. (15) is

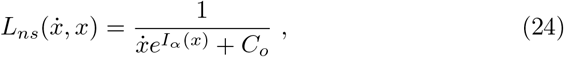

where the constant *C_o_* replaces the function *G*(*x*) in Eq. (6); the constant can have any real value, and it is not required to determine its value. This non-standard Lagrangian has no restrictions or limitations and it exists for any differentiable *α*(*x*) regardless of the forms of *β*(*x*) and *γ*(*x*). Therefore, 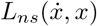 will be used to find NSLs for the population dynamics models (see Section 3). Moreover, the validity of one of the conditions given by Eqs (9) and (10) must be verified for all the models to determine whether any of these models allows for the NSL given by Eq. (6).

## 3 Population dynamics models and methods

### 3.1 Selected models

We consider the following population dynamics models: the Lotka-Volterra [33,34], Verhulst [35], Gompertz [36] and Host-Parasite [37] models that describe two interacting (preys and predators) species. However, the SIR model [38] describes the spread of a disease in a given population.

The variables *w*_1_(*t*) and *w*_2_(*t*) in the first four models of Table 1 represent prey and predators of the interacting species, respectively. Moreover, the time derivatives 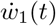 and 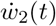 describe changes of these species in time. The interaction between the species in each model is given by the coefficients *a, A, b, B f*_1_, *f*_2_, *m*_1_ and *m*_2_, which are real and constant. However, in the SIR model, the variables *w*_1_(*t*) and *w*_2_(*t*) describe susceptible and infectious populations, with *a* and *b* being the recovery and infection rates, respectively.

**Table 1.** Biological models and their equations of motion.

| Population models | Equations of Motion |
| --- | --- |
| Lotka-Volterra Model | $\dot{w}_1 = w_1(a + bw_2)$<br>$\dot{w}_2 = w_2(A + Bw_1)$ |
| Verhulst Model | $\dot{w}_1 = w_1(A + Bw_1 + f_1w_2)$<br>$\dot{w}_2 = w_2(a + bw_2 + f_2w_1)$ |
| Gompertz Model | $\dot{w}_1 = w_1(A \log\left(\frac{w_1}{m_1}\right) + Bw_2)$<br>$\dot{w}_2 = w_2(a \log\left(\frac{w_2}{m_2}\right) + bw_1)$ |
| Host-Parasite Model | $\dot{w}_1 = w_1(a - bw_2)$<br>$\dot{w}_2 = w_2(A - B\frac{w_2}{w_1})$ |
| SIR Model | $\dot{w}_1 = -bw_1w_2$<br>$\dot{w}_2 = bw_1w_2 - aw_2$ |

Among the models given in Table 1, the Lotka-Volterra, Verhulst and Gompertz models are symmetric, and the remaining two models are asymmetric, where being symmetric means that the dependent variables can be replaced by each other, and the same can be done with the coefficients. Obviously, this cannot be done for the asymmetric models.

### 3.2 Methods to construct non-standard Lagrangians

According to Table 1, each model is described by a coupled first-order nonlinear ODEs that can be cast into a second-order nonlinear ODE of the following form

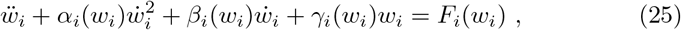

where *i* = 1 and 2. This equation is of the same form as that given by Eq. (5). The non-standard Lagrangian for this equation is

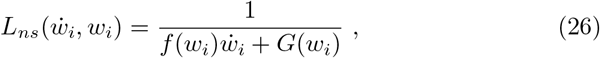

with the functions *f*(*w_i_*) and *G*(*w_i_*) being given by Eqs (7) and (8), respectively. Thus, for each considered model and for each variable in this model, we determine these functions and obtain 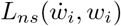.

Then, we verify the validity of one of the conditions given by Eqs (9) and (10). Since these conditions are equivalent, we take only one of them and select the condition on *γ*(*w_i_*) (see Eq. 10). The explicit form of this condition used in our calculations is

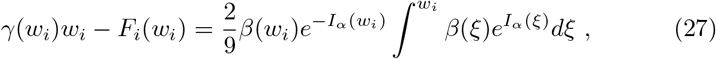

This condition is used to verify whether the derived NSL (see Eq. 6) can be constructed for Eq (5) or not (see Proposition 1). Moreover, the validity of this condition determines uniquely that the equation of motion can be converted into a linear second-order ODE, whose solutions are easy to find (see Proposition 2).

As shown in Section 2.3, another method to construct NSLs is to cast Eq. (25) into the form

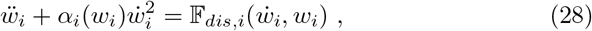

where

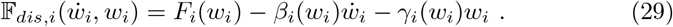

Then, according to Eq. (18), the NSL for Eq. (28) is given by

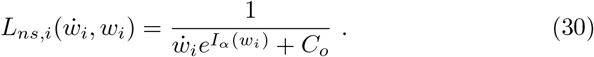

This non-standard Lagrangian is not constrained by any conditions and it can be derived for all considered population dynamics models. If this NSL is substituted into the following EL equation

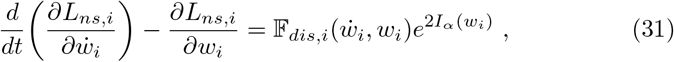

then, the equation of motion for the considered model is obtained. In the following, this method is used to construct the NSLs for all population dynamics models considered in this paper.

## 4 Models and their non-standard Lagrangians

### 4.1 Lotka-Volterra Model

The model is symmetric and it is represented mathematically by a system of coupled nonlinear first-order ODEs given in Table 1. We cast the first-order ODEs into the second-order ODEs of the form given by Eq. (28), and obtain

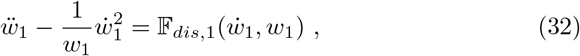

where

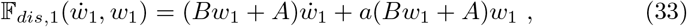

and

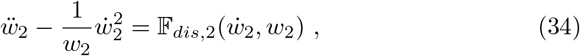

with

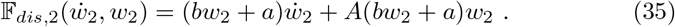

Using Eq. (7), the factors *e*^*I_α_*(*w_i_*)^ for both models can be calculated, and the obtained results are substituted into Eq. (30) to give

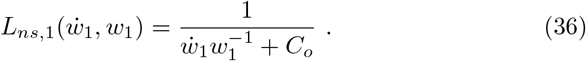

and

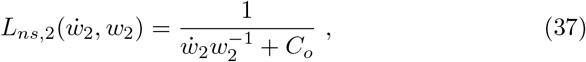

which are the NSLs for the Lotka-Volterra model. It is easy to verify that by substituting them into the E–L equation given by Eq. (31), the evolution equations describing the Lotka-Volterra model are obtained (see Eqs 32 and 34).

### 4.2 Verhulst Model

The system of coupled nonlinear ODEs given in Table 1 shows that the model is symmetric. The second-order equations for the dynamical variables of this model are:

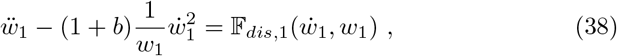

where

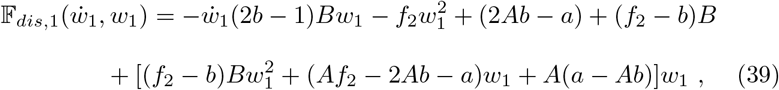

and

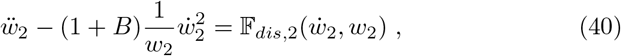

with

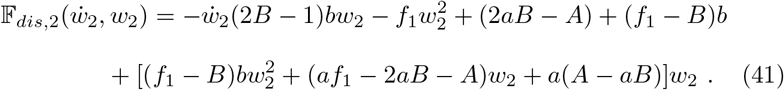

The non-standard Lagrangians for the evolution equations describing this model are:

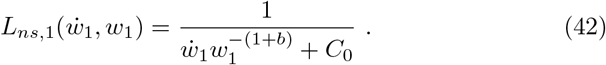

and

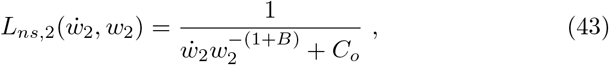

The derived NSLs give the original evolution equations for the model (see Eqs 38 and 40) after they are substituted into the E–L equation given by Eq. (31).

### 4.3 Gompertz Model

The mathematical representation of this model given by the coupled and nonlinear ODEs in Table 1 show that the model is symmetric.

The equation describing the time evolution of each model variable is given as

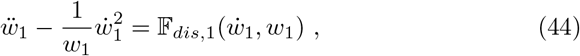

where

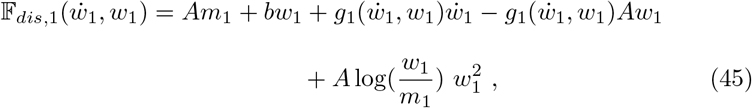

and

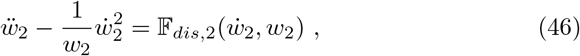

with

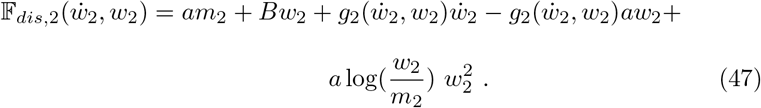

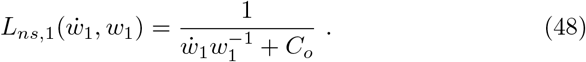

and

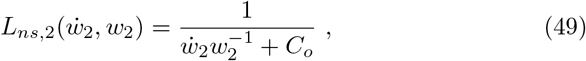

To obtain the original evolution equations given by Eqs 44 and 46) for this model, it is necessary to substitute the derived NSLs into the E–L equation (see Eq. 31).

### 4.4 Host-Parasite Model

This model describes the interaction between a host and its parasite. The model takes into account the non-linear effects of the host population size on the growth rate of the parasite population [22]. The system of coupled nonlinear ODEs (see Table 1) is asymmetric in the dependent variables *w*_1_ and *w*_2_. The time evolution equations for these variables are:

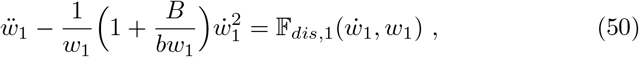

where

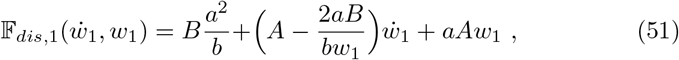

and

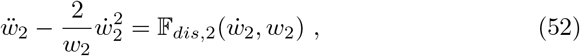

with

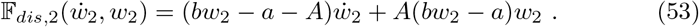

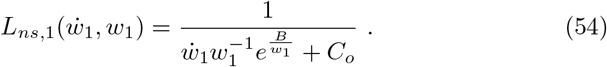

and

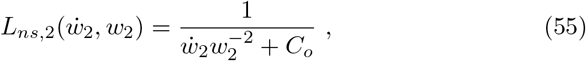

Then, the original evolution equations for this model (see Eqs 50 and 52) is obtained by substituting the derived NSLs into the E–L equation (see Eq. 31).

### 4.5 SIR Model

The equation describing the time evolution of each model variable is given as

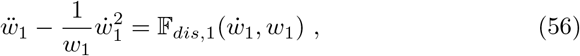

where

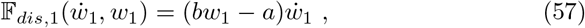

and

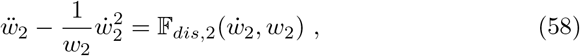

with

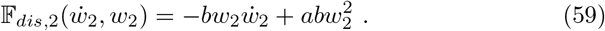

Using Eq. (7), the factors *e*^*I_α_*(*w_i_*)^ for both models can be calculated, and the obtained results are substituted into Eq. (30) to give

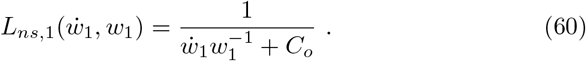

and

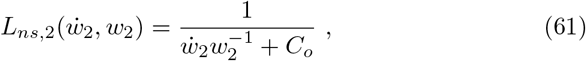

The SIR model is asymmetric and the derived NSLs give the original evolution equations for this model (see Eqs 56 and 58) after the NSLs are substituted into the E–L equation given by Eq. (31).

## 5 Null Lagrangians for the population models

In our previous work [32], we showed how to construct standard Lagrangians for the population dynamics models. Moreover, in this paper, we constructed the NSLs for the same models. However, there is another family of Lagrangians called null Lagrangians (NLs), which make the E–L equation identically zero and are given as the total derivative of a scalar function [39]; the latter is called here a gauge function. The NLs were extensively studied in mathematics (e.g., [39–44]) and recently in physics [45–49], but to the best of our knowledge, NSLs have not yet been introduced to biology and, specifically, to its population dynamics. Therefore, in the following, we present the first applications of NLs to biology and its population dynamics.

Recent studies of null Lagrangians demonstrated that there is a different condition that is obeyed by NLs and that this condition plays the same role for NLs as the E–L equation plays for standard and non-standard Lagrangians [48,49]. The condition can be written as

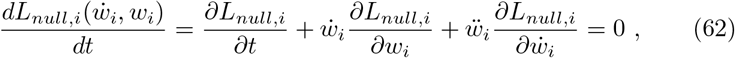

and it shows that the substitution of any NL into Eq. (62) results in an equation of motion. However, the resulting equations of motion may be limited because their coefficients are required to obey relationships that are different for different equations of motion [48,49]. The previous work also demonstrated that an inverse of any null Lagrangian generates a non-standard Lagrangian, whose substitution into the E–L equation gives a new equation of motion. However, the reverse is not always true, which means that not all NSLs have their corresponding NLs because of the so-called null condition that must be satisfied [48,49].

For any non-standard Lagrangian of the form

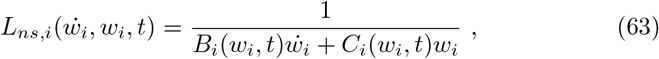

the null condition [48,49] is

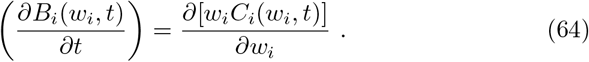

Comparison of Eq. (63) to Eq. (30) shows that *B_i_*(*w_i_*) = *exp*[*I_α_*(*w_i_*)] and *w_i_ C_i_*(*w_i_,t*) = *C_o_*, which means that the denominator of Eq. (30) satisfies the null condition and, therefore, the null Lagrangians for all the considered population dynamics systems are of the form

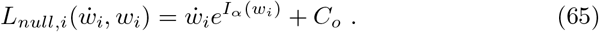

It is easy to verify that substitution of 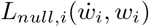 into the E–L equation gives identically zero.

Now, to derive the equation of motion given by Eq. (31), the null condition of Eq. (62) must be modified to account for the dissipative force. Then, the null condition is

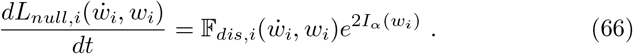

Substitution of 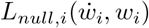 into this equation gives the required equation of motion (see Eq. 31). In the following, we use Eq. (65) to find the NLs for all the population dynamics models considered in this paper.

Another important characteristic of NLs is the fact that they can always be expressed as the total derivative of a scalar function [39], which has been called a gauge function [45–49]. Thus, we may write

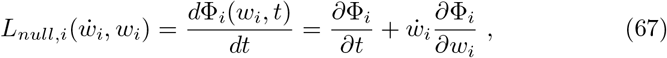

and use it to determine the gauge functions Φ_*i*_(*w_i_, t*) for all derived null Lagrangian 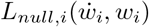.

The results of our derivations of the null Lagrangians (see Eq. 65) and their corresponding gauge functions (see Eq. 67) are presented in Table 2. To the best of our knowledge, these are the first null Lagrangians and gauge functions ever presented for any biological systems, especially for the population dynamics models.

**Table 2.** Null Lagrangians and their gauge functions for the population models

| Population models | Null Lagrangians | Gauge functions |
| --- | --- | --- |
| Lotka-Volterra | $L_{null,1} = \dot{w}_1 w_1^{-1} + C_0$<br>$L_{null,2} = \dot{w}_2 w_2^{-1} + C_0$ | $\Phi_1 = \ln \vert w_1\vert + C_0 t$<br>$\Phi_2 = \ln \vert w_2\vert + C_0 t$ |
| Verhulst | $L_{null,1} = \dot{w}_1 w_1^{-(1+b)} + C_0$<br>$L_{null,2} = \dot{w}_2 w_2^{-(1+B)} + C_0$ | $\Phi_1 = -b^{-1} w_1^{-b} + C_0 t$<br>$\Phi_2 = -B^{-1} w_1^{-B} + C_0 t$ |
| Gompertz | $L_{null,1} = \dot{w}_1 w_1^{-1} + C_0$<br>$L_{null,2} = \dot{w}_2 w_2^{-1} + C_0$ | $\Phi_1 = \ln \vert w_1\vert + C_0 t$<br>$\Phi_2 = \ln \vert w_2\vert + C_0 t$ |
| Host-Parasite | $L_{null,1} = \dot{w}_1 w_1^{-1} e^{\frac{B}{w_1}} + C_0$<br>$L_{null,2} = \dot{w}_2 w_2^{-2} + C_0$ | $\Phi_1 = -Ei(\frac{B}{w_1}) + C_0 t$<br>$\Phi_2 = -w_2^{-1} + C_0 t$ |
| SIR | $L_{null,1} = \dot{w}_1 w_1^{-1} + C_0$<br>$L_{null,2} = \dot{w}_2 w_2^{-1} + C_0$ | $\Phi_1 = \ln \vert w_1\vert + C_0 t$<br>$\Phi_2 = \ln \vert w_2\vert + C_0 t$ |

## 6 Discussion and comparison to the previous work

Different methods to construct standard and nonstandard Lagrangians have been developed and applied to various dynamical systems. In the constructed standard Lagrangians, the presence of the kinetic and potential energy-like terms is evident [1–15,32]. However, the previously constructed NSLs have different forms, in which such energy-like terms cannot be easily recognized [12–31]. Some previously obtained NSLs were of the forms given by either Eq. (1) or Eq. (3) [12–15,21,26], but other NSLs had significantly different forms [17–19,22–25]. It is important to note that the NSLs obtained using the Jacobi Last Multiplier method developed by Nucci and Leach (e.g., [17–19]), and those obtained by El-Nabulsi (e.g., [22–24]) using a different method, have been applied to many various dynamical systems in applied mathematics, physics, and astronomy.

In the first attempts to obtain Lagrangians for selected biological systems, the Lagrangians were found by guessing their forms for selected population dynamics models [27–29]. Later, those Lagrangians were formally derived by Nucci and Tamizhmani [30], using the method of Jacobi Last Multiplier; see also [31] for other applications of this method to biological systems. Interesting recent work was done by Carinena and Fernandez-Nunez [50], who considered systems of first-order equations and derived the NSLs that are linear, or more generally affine, in velocities using the method of Jacobi Last Multiplier. Among the applications of their results to different dynamical systems, they also included several population dynamics models [50].

In our previous work [32], we derived the first standard Lagrangians for five population dynamics models and discussed physical and biological implications of these Lagrangians for the models. In this paper, we derived the NSLs by using a method that significantly modified the one previously developed [12,20]. The results obtained in this paper are now compared to those previously obtained by Nucci and Tamizhmani [30], as they are the most relevant to our work. The comparison shows that there are some advantages of using the method of Jacobi Last Multiplier, such as the method does not require introducing forcing functions and gives directly the same Lagrangians as those found earlier in [27–29]. However, the advantage of the method developed in this paper is that the derived NSLs are directly related to their null Lagrangians (NLs) and gauge functions (GFs).

Having obtained the NSLs for the considered population dynamics models, we use them to find their corresponding NLs and then to derive the gauge functions (see Table 2); to the best of our knowledge, these NLs and GFs are the first obtained for biological systems, and specifically, for the population dynamics models. An interesting result is that the NLs and their gauge functions are identical for the Lotka-Volterra, Gompertz, and SIR models, which are caused by the same forms of NSLs for these three models. It must also be noted that the gauge functions are given by the logarithmic functions, which is a novel form among the previously obtained gauge functions (e.g., [45–49]). In the approach presented in this paper, the main differences between the models are shown by the introduced dissipative functions.

Now, the NSLs, NLs and GFs for the Verhulst and Host-Parasite models have significantly different forms when compared to the other three models (see Table 2). Moreover, there are also differences in their NSLs and NLs. A new result is the form of the gauge function Φ_1_ for the Host-Parasite model as this function is given by the exponential function *Ei*(*B*/*w*_1_), which also appeared in the standard Lagrangian found for this system in [32]; this implies that there exist some relationships between standard, non-standard and null Lagrangians, and that the gauge function plays an important role in such relationships; further exploration of such relationships will be done in a separate paper.

## 7 Conclusions

A new method to derive non-standard Lagrangians for second-order nonlinear differential equations with damping is developed and applied to the Lotka-Volterra, Verhulst, Gompertz, Host-Parasite, and SIR population dynamics models. For the considered models, the method shows some limitations, which are explored and it is demonstrated that these limitations do not exist for the models whose equations of motion can be converted into linear ones.

The obtained non-standard Lagrangians are different than those previously obtained for the same models [30], and the main difference is that the previously used Jacobi Last Multiplier method does not require introducing dissipative forcing functions, which were defined in this paper. However, the advantage of the method developed in this paper is that it allows us to use the derived non-standard Lagrangians to obtain first null Lagrangians and their gauge functions for the population dynamics models.

By following the recent work [48,49], the presented results also demonstrate how the derived null Lagrangians and gauge functions can be used to obtain the equations of motion for the considered population dynamics models. Our approach to solving the inverse calculus of variation problem and deriving non-standard and null Lagrangians is applied to the models of population dynamics. However, the presented results show that the method can be easily extended to other biological or physical dynamical systems whose equations of motion are known.

## Acknowledgement

We thank Rupam Das and Lesley Vestal for discussions of null Lagrangians and gauge functions and their applications to different dynamical systems.

## Notes

### Competing Interest Statement

The authors have declared no competing interest.

## References

[1] J.L. Lagrange, Analytical Mechanics (Springer, Netherlands, 1997).

[2] H. Goldstein, C.P. Poole, J.L. Safko, Classical Mechanics, 3rd Edition (Addison-Wesley, San Francisco, CA, 2002).

[3] J.V. José, E.J. Saletan, Classical Dynamics, A Contemporary Approach, (Cambridge Univ. Press, Cambridge, 2002).

[4] Lopuszanski, J., The Inverse Variational Problems in Classical Mechanics (World Scientific, Singapore, 1999).

[5] N.A. Daughty, Lagrangian Interactions (Addison-Wesley Publ. Comp. Inc. Sydney, 1990).

[6] V.I. Arnold, Mathematical Methods of Classical Mechanics (Springer, New York, NY, USA, 1978).

[7] P.J. Olver, Applications of Lie Groups to Differential Equations (Springer-Verlag, New York, 1993).

[8] H. Helmholtz, J. Reine Angew Math. 100, 213, 1887.

[9] J. Douglas, Trans. Am. Math. Soc. 50, 71–128, 1941.

[10] S. A. Hojman, J. Phys. A: Math. Gen. 17, 2399–2412, 1984.

[11] S.A. Hojman, J. Phys. A: Math. Gen. 25, L291 – L297, 1992.

[12] Z.E. Musielak, J. Phys. A Math. Theor. 41, 055205, 2008.

[13] J.L. Ciéslinski and T. Nikiciuk, J. Phys. A Math. Theor. 43, 175205, 2010.

[14] Z.E. Musielak, N. Davachi and M. Rosario-Franco, Mathematics. 8, 379, 2020.

[15] Z.E. Musielak, N. Davachi and M. Rosario-Franco, J. Appl. Math. ID 3170130 (11 pages), 2020.

[16] A.I. Alekseev and B.A. Arbuzov, Theor. Math. Phys. 59, 372–378, 1984.

[17] M.C. Nucci and P.G.L. Leach, J. Math. Phys. 48, 123510, 2007.

[18] M.C. Nucci and P.G.L. Leach, J. Math. Phys. 49, 073517, 2008

[19] M.C. Nucci and P.G.L. Leach, Phys. Scripta. 78, 065011, 2008

[20] Z.E. Musielak, Chaos, Solitons Fractals, 42, 2640, 2009.

[21] A. Saha and B. Talukdar, Rep. Math. Phys. 73, 299–309, 2014.

[22] R.A. El-Nabulsi, App. Math. Lett. 24, 1647, 2011.

[23] R.A. El-Nabulsi, Qual. Theory Dyn. Syst. 12, 273, 2013.

[24] R.A. El-Nabulsi, Int. J. Theor. Phys. 56, 1159, 2017.

[25] F.E. Udwadia, H. Cho, J. Appl. Mech. 80, 041023, 2013.

[26] N. Davachi and Z.E. Musielak, J. Undergrad. Rep. Phys. 29, 100004, 2019.

[27] E.H. Kerner, Bulletin of Mathematical Biophysics. 26, 333–349, 1964.

[28] S.L. Trubatch and A. Franco, J. Theor. Biology. 48, 299–324, 1974.

[29] G.H. Paine, Bulletin of Mathematical Biology. 44, 749–760, 1982.

[30] M.C. Nucci and K.M. Tamizhmani, J. Nonlinear Math. Phys. 19, 12500021, 2012.

[31] M.C. Nucci and G. Sanchini, Symmetry, 7, 1613–1632, 2015.

[32] D.T. Pham and Z.E. Musielak, Phys. Scripta. submitted, 2022; arXiv:2203.13138v1 [q-bio-PE] 24 March 2022.

[33] A.J. Lotka, Elements of Physical Biology (Baltimore, 1925).

[34] V. Volterra, Nature. 18, 1–42, 1926.

[35] P.F. Verhulst, Correspondance mathématique et physique, 10, 113–21, 1838.

[36] B. Gompertz, Phil Trans Roy Soc. 27, 513–85, 1825.

[37] V.P. Collins, R.K. Loeffler, H. Tivey, Am J Roentgenol Radium Ther Nuc Med. 78, 988–1000, 1956.

[38] W.O. Kermack and A.G. McKendrick, Proc. Roy. Soc. Lond. A. 115, 700–721, 1927.

[39] P.J. Olver, Applications of Lie Groups to Differential Equations (Springer-Verlag, New York, 1993)

[40] P.J. Olver and J. Sivaloganathan, Nonlinearity. 1, 389, 1989.

[41] M. Crampin and D.J. Saunders, Diff. Geom. and its Appl. 22, 131, 2005.

[42] R. Vitolo, Diff. Geom. and its Appl. 10, 293, 1999.

[43] D. Krupka and J. Musilova, Diff. Geom. and its Appl. 9, 225, 1998.

[44] D. Krupka. O. Krupkova, and D. Saunders, Int. J. Geom. Meth. Mod. Phys. 7, 631, 2010.

[45] Z. E. Musielak and T. B. Watson, Phys. Let. A. 384, 126642, 2020.

[46] Z.E. Musielak and T.B. Watson, Phys. Let. A. 384, 126838, 2020.

[47] L.C. Vestal, Z.E. Musielak, Physics. 3, 449, 2021.

[48] R. Das and Z.E. Musielak, Phys. Scripta. 97, 125213 (12 pages), 2022.

[49] R. Das and Z.E. Musielak, Phys. Scripta. submitted, 2022; arXiv:2210.09105v1 [math-ph] 17 Oct 2022.

[50] J.F. Carinena and J. Fernandez-Nunez, Symmetry. 14, 2520, 2022.

